# Plant specialised metabolites modulate the molecular signatures of host-bacteria and bacteria-bacteria interactions

**DOI:** 10.1101/2025.06.06.658260

**Authors:** Liza Rouyer, Claude Becker, Niklas Schandry

## Abstract

Plants participate in intricate interactions with a multitude of microorganisms, many of which also influence each other. This holobiont is situated in a chemical soil environment that is defined, in parts, by the specialised metabolite legacy of proximal and preceding organisms, including other plants. Here, we investigated the influence of external plant-derived specialised metabolites on the interactions among root-associated bacterial strains, and between these strains and a plant host. Using benzoxazinoids and their derivatives as a model in both simplified pairwise experiments and more complex multi-organism analyses, we show that these chemicals can modulate bacteria-bacteria, as well as bacteria-plant interactions. While the chemical environment alone had little effect on the plant at the molecular level, it differentially affected plant chemical defences, immunity, and sugar transport when combined with single-isolate or micro-community inoculums. Our study underlines the importance of the chemical environment in modulating organismic interactions and illustrates the value of combining reduced-complexity, bottom-up reconstruction approaches with top-down holobiont profiling.

**SIGNIFICANCE:** Many plant species secrete specialized metabolites into the soil, where they can have a long-lasting effect on subsequent plant generations and their associated microbiomes. Understanding the effect of this chemical environment on soil- and plant-associated microbiomes is crucial to determine the impact of soil legacy on host plants, for example in the context of crop rotations. Here, we report that the interactions among root-associated microbes are modulated by specialized metabolites of the benzoxazinoid family, which are prominent metabolites in many grasses. We further show that the chemical environment can inhibit the defence capacity of the plant towards colonizing bacteria, and that more complex bacterial communities are able to mitigate these effects. Our work highlights the importance of deconstructing bacterial communities and the chemical environment to gain insights into the fine-tuned molecular mechanisms that determine the outcome of complex organismic interactions.

## INTRODUCTION

Microbiomes are complex ecosystems that can vary depending on the plant species, environmental conditions, and the host’s life cycle, implying a tight regulation of microbiome composition by the host (Fitzpatrick et al., 2020). The plant immune system is tailored to recognise conserved microbial molecules via pattern-recognition receptors (PRRs), which leads to the activation of pattern-triggered immunity (PTI) as the first line of defence (Jones et al., 2024). As pathogens evolved effector molecules that interfere with host responses, plants evolved effector-triggered immunity (ETI) as a second line of defence, by which Nucleotide-binding Leucine-rich Repeat (NLR) receptors sense microbial effectors and trigger various defence mechanisms (Jones et al., 2024). Although tailored to fend off pathogens, plant immunity also acts towards non-pathogenic microbes (Fitzpatrick et al., 2020; Teixeira et al., 2019). To colonise a host, commensal bacteria consequently need to escape plant immunity by suppressing plant responses or evading host recognition (Teixeira et al., 2019). For example, rhizosphere bacteria of the Xanthomonadales family can modulate the immune system of *Arabidopsis thaliana*, leading to reduced immune responses and a competitive advantage in root colonisation (Ordon et al., 2025).

Moreover, plants can recruit and shape their microbiome through the exudation of up to 40% of their assimilated carbon in the form of various organic molecules, mostly sugars (An et al., 2019; Canarini et al., 2019; Kim et al., 2021; Sasse et al., 2018). The SWEET (<u>S</u>ugars <u>W</u>ill <u>E</u>ventually be <u>E</u>xported <u>T</u>ransporters) sugar uniporters, are differentially expressed along the root axis of Arabidopsis, allowing for spatial organisation of specific microbial communities (Chen et al., 2010; Loo et al., 2024). Plant sugar transport proteins (STPs) are responsible for sugar import, lowering sugar levels in the apoplast, and are involved in defence responses by depriving pathogens of nutrients (Lemonnier et al., 2014), which highlights the importance of sugar transport in microbiome and pathobiome establishment.

Plant-derived specialised metabolites also play a role in modulating the microbiome and have been increasingly recognised as the primary route for this inter-kingdom communication (Korenblum et al., 2022; Pang et al., 2021; Weston and Mathesius, 2013; Zaynab et al., 2018). For example, glucosinolates (GLS) from Brassicaceae attract specific microbes, which in turn influence GLS production (DeWolf et al., 2023); strigolactone content is increased in phosphorus-deficient conditions to recruit arbuscular mycorrhiza to the roots (Yoneyama et al., 2007); and coumarins recruit specific root-associated bacteria, leading to induced systemic resistance (ISR) in the plant (Stassen et al., 2021). Similarly, the benzoxazinoids (BXs) DI(M)BOA (2,4-dihydroxy-(methoxy)-1,4-benzoxazin-3-one), which can be found in many grass species, are exuded through the roots and have an important and long-lasting impact on soil microbiomes (Cotton et al., 2019; Gfeller et al., 2023; Hu et al., 2018; Janse van Rensburg et al., 2025; Stengele et al., 2024; Thoenen et al., 2023). Upon release into the soil, DIBOA is rapidly converted to BOA (2-benzoxazolinone) and ultimately to APO (2-amino-3H-phenoxazin-3-one), which exhibits high phytotoxicity and antibacterial properties, remaining detectable in soil for years (Anzai et al., 1960; Macías et al., 2004; Schandry et al., 2021; Venturelli et al., 2016). Such long-lasting effects of plant specialised metabolites in soil are of particular importance in agricultural contexts, as the legacy left by previous crops will impact the growth of the next generation (Gfeller et al., 2023; Janse van Rensburg et al., 2025; Semchenko et al., 2022; Stengele et al., 2024; Sun et al., 2014). Interlinked with trans-generational effects, the composition of the soil microbiome is driven by plant specialised metabolites and is a major player in soil legacy (Bakker et al., 2018; Hu et al., 2018; Janse van Rensburg et al., 2025; Steinauer et al., 2023). However, the basic mechanisms through which plant specialised metabolites act on the microbiome composition and function remain unclear, which is particularly relevant if one considers that stable metabolites remain active in the soil even after the plant that produced them has disappeared.

In recent years, many approaches have been used to try and break down the complexity of plant-microbe-soil interactions. On the one hand, bottom-up approaches, which apply the design of communities composed of members with known characteristics, allow for a tight control of each member of the community. They have led to major advances in the understanding of commensal-host homeostasis and protective capabilities of bacterial strains adapted to abiotic stress (Ma et al., 2021; Schmitz et al., 2022; Voges et al., 2019). Top-down approaches, on the other hand, take existing communities and aim at understanding them as a whole, for example via the introduction of perturbations and the identification of essential members (Jing et al., 2024). They have led to significant advances in understanding the role of plant hormones in plant-bacteria communication (Conway et al., 2022; Finkel et al., 2020), on the extent of plant responses to bacterial inoculums (Keppler et al., 2025), and on the regulation of plant stress responses by root commensals (Hou et al., 2021).

In this study, we set out to expand reductionist bottom-up approaches to understand the impact of metabolite legacy on microbiome assembly and plant responses. We used benzoxazinoids (BX) as a representative of well-studied legacy-conferring plant specialised metabolites.

After initially establishing that binary bacteria-bacteria interactions are modulated by the chemical environment *in vitro*, we used a tightly controlled experimental setup to investigate ternary bacteria-host-chemical interactions and generated a top-down meta-transcriptomics dataset to investigate the interactions across different levels of complexity. The presence of benzoxazinoids lead to a dampening of the host’s hormonal signalling and a subsequent decrease of defence responses. Intriguingly, the inoculation with more complex communities attenuated this effect of the chemical environment. In summary, our study highlights the finely tuned responses of bacteria and plants to the chemical environment and illustrates the importance of deconstructing and reconstructing complex interactions, looking beyond the biotic environment to understand multi-organismic relationships.

## RESULTS

### Inhibition of bacterial rhizosphere isolates is modulated by plant specialised metabolites

To investigate the effects of plant-derived chemicals on bacterial interactions, we first tested the impact of chemical treatment on pairwise bacterial interactions, in absence of a host. We used a curated collection of bacterial isolates from the roots of Arabidopsis and *Lotus japonicus* (Bai et al., 2015; Finkel et al., 2020; Wippel et al., 2021), comprising 73 isolates and spanning six bacterial orders and three phyla, the identity of which was confirmed via whole genome sequencing (Figure 1a).

**Figure 1.**
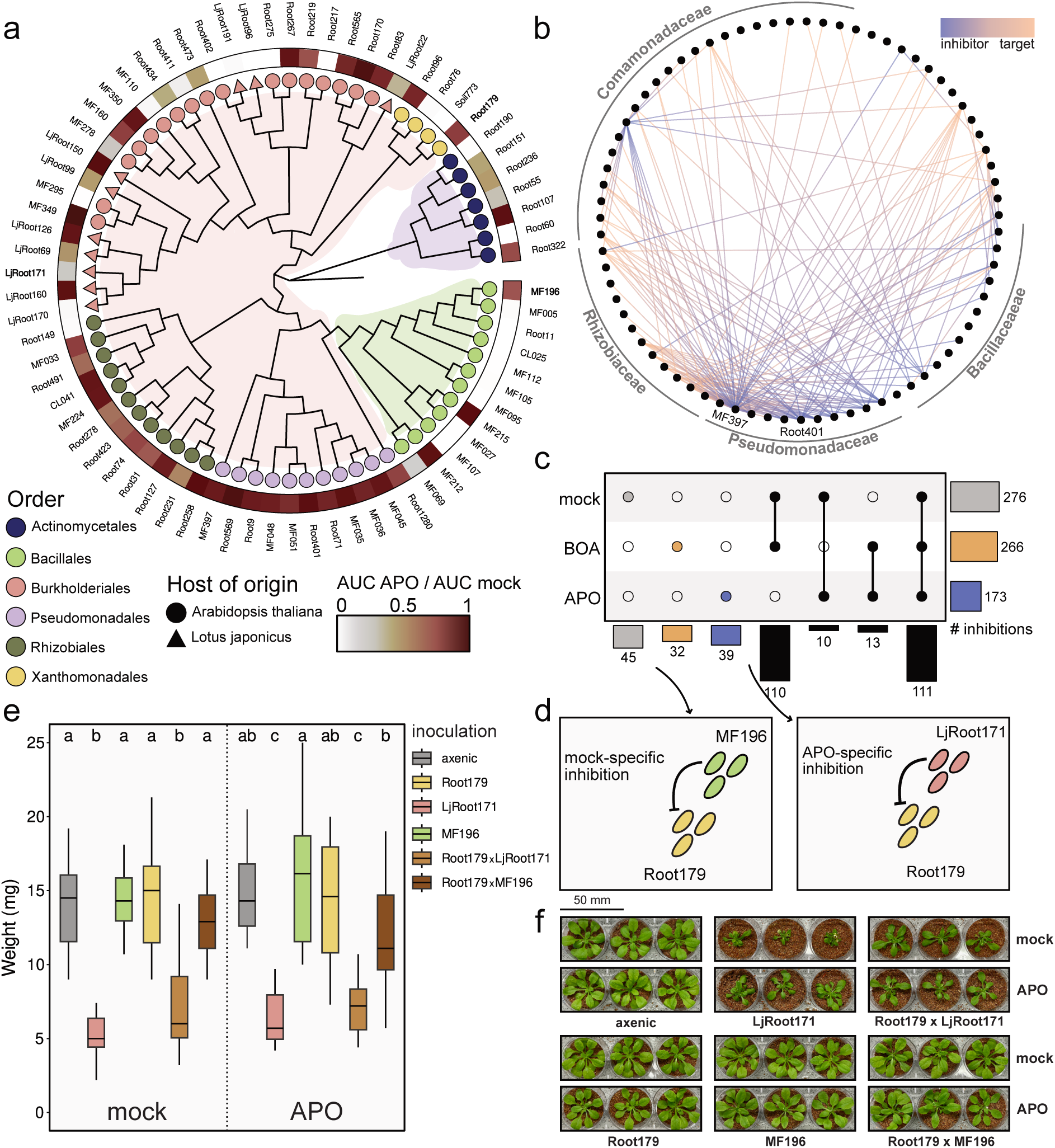
Bacteria-bacteria interactions are modulated by plant specialised metabolites. **a**. Phylogenetic tree of the bacterial isolates used in this study. Node colour indicates order, the outer ring shows the ratio between the area under the curve (AUC) of isolates grown in APO-containing and those grown in control medium. **b**. Network representation of *in vitro* binary interactions between 73 isolates. Nodes indicate isolates, and inhibitions are drawn as edges between nodes; colour gradient of the edges (purple → orange) indicates direction of the inhibition (inhibitor → inhibited). **c**. Upset plot showing the number of inhibitions in each chemical context (rows) and the ones in different combinations of conditions (columns). **d**. Specific interactions of the focus isolates used in this study. **e**. Rosette weight of Arabidopsis, axenic or inoculated. Different letters indicate significant statistical difference (p<0.05; ANOVA followed by a TukeyHSD test. **f**. Representative pictures of Arabidopsis Col-0 plants inoculated with single isolates and micro-communities, in mock conditions.

Testing all pairwise combinations for growth inhibition (5,625 interactions in two biological replicates), we observed 276 inhibitory interactions (5% of all pairs; Figure 1b). Of these, 63% were caused by the genus *Pseudomonas* and 25% by *Bacillus*, even though these genera accounted only for respectively 13% and 16% of the tested isolates. These results are in line with the findings of Helfrich et al. (2018), who surveyed over 200 Arabidopsis leaf isolates for growth interference and found that *Bacillus* and *Pseudomonas* dominated the inhibition network. While most of the inhibitory strains inhibited the growth of only a few isolates, suggesting specific interactions, ten isolates accounted for 75% of the inhibitions, with each one inhibiting more than ten other isolates. Most notably, *Pseudomonas* MF397 inhibited over two-thirds of all isolates, suggesting that this strain produced several antibiotics that targeted most of the surveyed genera. *Pseudomonas* Root401, which had previously been reported to have strong inhibitory capacity due to its ability to produce both pyoverdine and the antimicrobial DAPG (Getzke et al., 2023), served as an internal control and was among the most suppressive isolates, inhibiting 11 others (Figure 1b).

The natural hosts of the isolates of our collection, Arabidopsis and Lotus, do not produce benzoxazinoids, therefore leading to isolates naive to these chemicals. To understand how bacteria-bacteria interactions are influenced by APO and BOA, we repeated the pairwise interaction assay, supplementing the growth medium with either of these metabolites. One third of all inhibitory interactions were affected by the chemical environment (Figure 1c): 12% of the interactions were inhibitory only in mock conditions, while 9% and 11% were specific to BOA and APO, respectively (Figure 1c). Only 3% of all inhibitory interactions occurred in both BOA and APO but not in mock conditions, suggesting that these two metabolites differentially influence inhibitory interactions between isolates. As some bacteria can convert BOA to the more potent APO, we next asked whether the differential inhibitory capacity of one third of all isolates was linked to their chemical phenotype. Of 73 isolates, 29 (40%) were capable of converting BOA to APO, while 55% (40/73) were inhibited by APO (Figure 1a, Supplementary Figure 1a). Interestingly, some isolates that were able to produce APO were also sensitive to it.

The capacity of a given isolate to inhibit other isolates in BOA-supplemented media was independent from its capacity to convert BOA to APO (Χ^2^ (df=1, n=209) = 0.05, p=0.8). Conversely, and contrary to our expectations, APO-sensitive isolates were not significantly inhibited by more isolates when APO was present (Χ^2^ (df=1, n=448) = 3.35, p=0.07). Taken together, these results indicate that the differences in the inhibitory capacity of APO- or BOA-containing media are linked neither to APO sensitivity of the inhibited isolates nor to the capacity of the inhibiting strains to produce APO. Given that APO has been reported as the long-term stable metabolic product in soil, we used only APO for subsequent assays in soil.

### Bacterial colonisation leads to transcriptional changes in the host

Benzoxazinoids modulate the microbiome composition (Kudjordjie et al., 2019), and this modulation cannot be explained by differential bacteria-bacteria interactions alone (Figure 1). To understand the influence of the chemical environment on differential microbial interactions, we next investigated how exogenous metabolites modulated bacteria-bacteria and plant-bacteria interactions at the molecular level in the presence of a host. We selected three bacterial isolates that had shown differential antagonistic behaviour *in vitro* (Figure 1d) and engaged in different types of interactions with Arabidopsis. The commensal *Rhodanobacter* Root179 was described to modulate Arabidopsis immunity, effectively evading detection from the host (Ordon et al., 2025); LjRoot171 (*Variovorax* sp.) reduced rosette weight of Arabidopsis (Figure 1e,f), while MF196 (*Bacillus* sp.) did not impact Arabidopsis growth (Figure 1e,f). We combined Root179 with MF196, which inhibited Root179 only in mock conditions, or LjRoot171, which inhibited Root179 only in the presence of APO (Figure 1d). We inoculated the selected isolates either individually or in micro-communities of two isolates on one-week-old Arabidopsis Col-0 seedlings and co-cultivated them for three weeks in presence or absence of APO (Supplementary Figure 1b). Interestingly, while LjRoot171 had a detrimental effect on the growth of Arabidopsis in both mock- and APO-treated environments, this effect was weak in APO (Figure 1e,f). After confirming, via counting colony-forming units (CFU), that all isolates had established themselves on the host roots at the end of the experiment (Supplementary Figure 1c), we extracted total RNA from the roots to carry out meta-transcriptomic sequencing. We mapped transcriptome reads to the Col-CEN reference genome (Naish et al., 2021) and the *de-novo* assembled bacterial genomes (Supplementary Figure 1d, see Material and Methods for details). A principal component analysis (PCA) based on all read counts (host and bacterial genes) showed a separation of samples by inoculum (Supplementary Figure 1e).

### APO interferes with bacterial defence abilities

In total, we recovered over 8 million bacterial reads, allowing us to follow transcriptomic responses of all three isolates *in planta*. We compared gene expression of each isolate when it was inoculated alone or in combination with one other isolate. We defined differentially expressed genes (DEGs) as those with an absolute log_2_(fold change) >1 and an FDR adjusted p-value < 0.05. For all pairwise inoculations, the presence of APO increased transcriptional responses compared to mock conditions. In the Root179xLjRoot171 combination, the presence of APO resulted in a 1.6-fold increase in the number of DEGs for both strains compared to mock (Figure 2a), while in the Root179xMF196 pair, the presence of APO lead to two times more DEGs in Root179 (Figure 2b).

**Figure 2.**
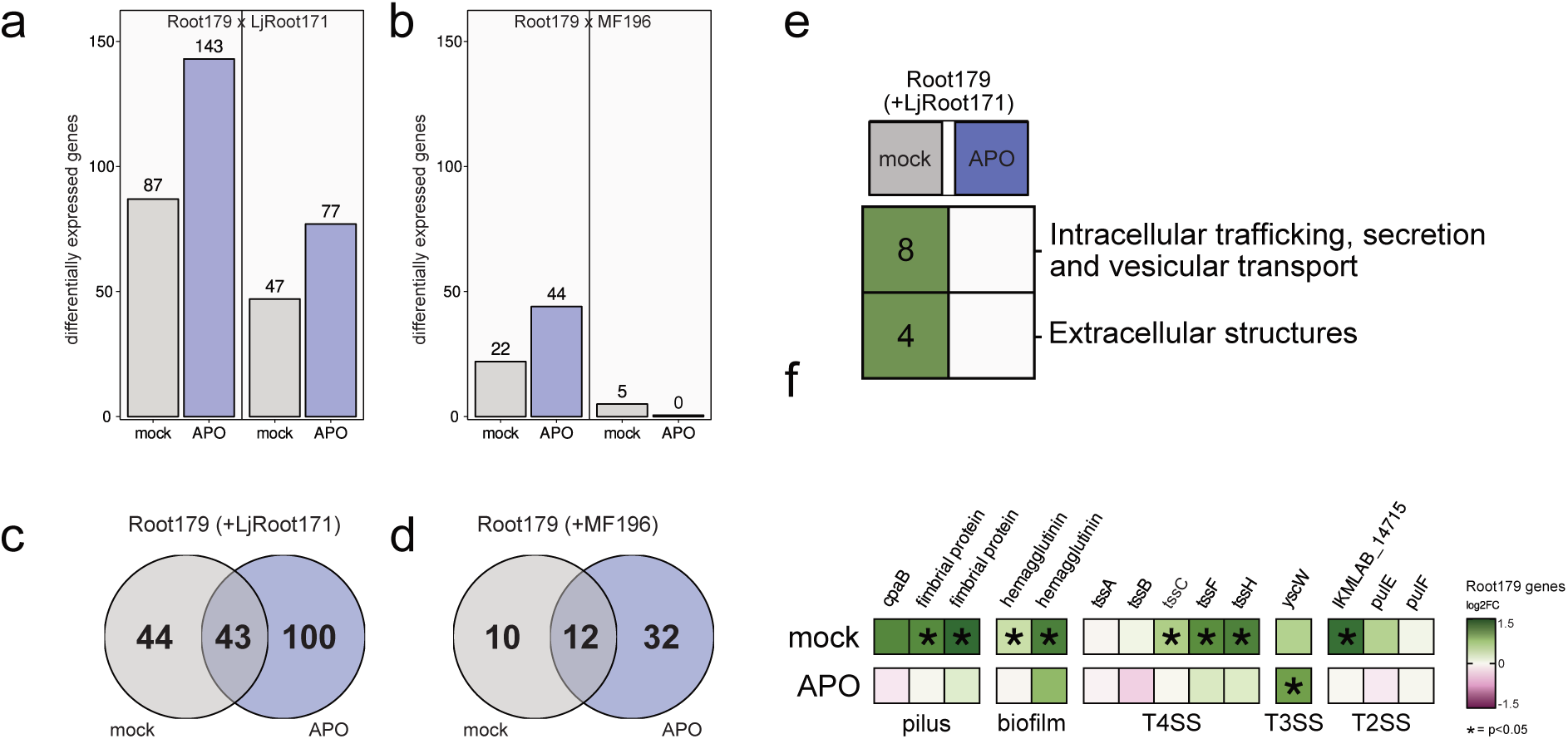
APO interferes with bacteria defence abilities. **a, b.** Number of differentially expressed genes (|L2FC| (absolute Log_2_(fold-change))> 1, FDR < 0.05) of Root179 and LjRoot171 (a) and Root179 and MF196 (b) when inoculated in micro-communities compared to their single inoculum counterpart. **c, d.** Root179 common and treatment-specific DEGs when co-inoculated with LjRoot171 (c) and MF196 (d). **e.** Cluster of orthologous gene (COG) enrichment analysis of Root179 genes showing differential expression when inoculated with LjRoot171. **f.** L2FC of Root179 genes related to bacterial competition.

In both tested pairs, Root179 responded more strongly to the presence of a second isolate than the respective co-inoculated isolate (Figure 2a,b, Supplementary 2a,b). The transcriptional response of pair Root179xLjRoot171 resulted in a total of 187 DEGs in Root179 (Figure 2c), that of the Root179xMF196 pair in 54 DEGs in Root179 (Figure 2d).

For both pairs, the *in vitro* assays showed that the inhibition of Root179 depended on the chemical environment, with LjRoot171 inhibiting Root179 only in presence of APO, and MF196 inhibiting Root179 only in absence of APO (Figure 1d). To gain functional insights into the specificity of this bacterial competition, we conducted a cluster of orthologous genes (COG) enrichment analysis, comparing DEGs of a single isolate to their micro-community inoculated counterpart. (Figure 2e, Supplementary Figure 2a). The two COG-terms “Intracellular trafficking, secretion and vesicular transport” and “Extracellular structures” were exclusive to Root179 genes upregulated in mock conditions in response to LjRoot171 (Figure 2e, Supplementary Figure 2c). This included genes of the type-2-secretion system T2SS (a pil-N domain containing protein, *pulE*, *pulF*), T3SS genes (*yscW*), and T4SS genes (*tssA*, *tssB*, *tssC*, *tssF*, *tssH*), pilus genes (*cpB*, two fimbrial proteins) and biofilm genes (two hemagglutinins) (Figure 2f). Taken together, this suggests that the competitiveness of Root179 towards LjRoot171 depends on the chemical environment and is suppressed in the presence of APO.

### Plant host responses are driven by bacterial inoculum rather than chemical treatment

While APO seemed to shape bacteria-bacteria competition *in planta*, consistent with our *in vitro* inhibition assays (Figure 1d, Figure 2e,f), we could not exclude an effect of the plant host onto these interactions. Our experimental set-up allowed an in-depth insight into transcriptional response of Arabidopsis facing selected isolates individually or in micro-communities, in the presence or absence of APO.

First, we focused on responses upon plant inoculation with a single isolate. The PCA of the 1,000 genes with the highest variance confirmed the previous observations of Ordon et al. (2025) that Root179 can evade host detection, as samples inoculated with Root179 were very similar to axenic samples (Figure 3a). This evasion was not influenced by APO, as the respective mock and APO samples both clustered with axenic samples. Samples inoculated with LjRoot171 showed the strongest response compared to axenic control; MF196-inoculated samples were spread between the axenic and LjRoot171-inoculated samples (Figure 3a). Overall, samples did not separate by chemical treatment but rather by bacterial inoculum. Interestingly, the response of Arabidopsis to APO seemed to depend on the presence of bacteria, as we detected only 5 DEGs in axenic plants in response to APO, compared to 16, 142 and 101 DEGs when inoculated with LjRoot171, Root179 and MF196, respectively (Figure 3b).

**Figure 3.**
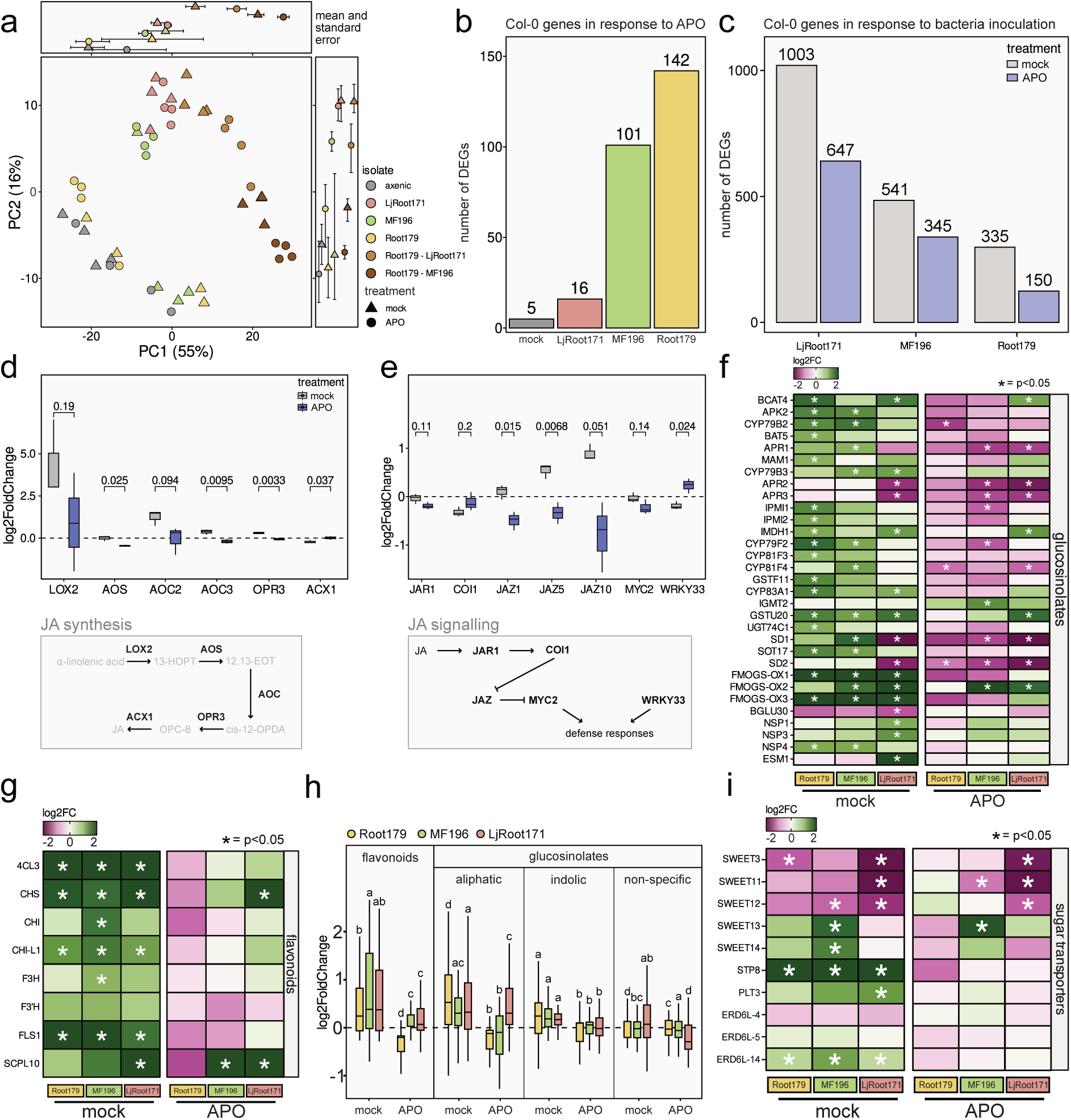
APO impairs Arabidopsis defence responses. **a.** Principal component analysis (PCA) on the top 1,000 Col-0 genes showing the highest variance between inoculated and axenic samples. Boxes on the top and right-side show mean and standard error for each condition, along PC1 and PC2. **b.** Number of Col-0 DEGs in response to APO treatment. **c.** Number of Col-0 DEGs in response to bacterial inoculation. **d, e.** L2FC of genes involved in jasmonic acid (JA) biosynthesis (d) and signalling (e), respectively (two-tailed Student’s t-test). The genes shown in the boxplots are highlighted in the pathways at the bottom of the plots. **f, g.** L2FC of genes involved in the biosynthesis of defence compounds from the flavonoid (f) and glucosinolate (g) families showing differential expression in at least one condition. The complete set of genes of these pathways and their differential expression can be found in Supplementary Figure 4. **h.** L2FC of all genes from the glucosinolates and flavonoids pathways detected in our analysis. Different letters indicate significant statistical difference (p<0.05; ANOVA followed by a TukeyHSD test). **i.** L2FC of genes encoding for sugar transporters that showed differential expression in at least one condition. Asterisks in e, f and h indicate significant differential expression (FDR < 0.05).

When comparing numbers of DEGs between mock and APO-treated samples for each bacterial inoculum, we found that APO reduced the effect of the bacteria on the transcriptional reprogramming of the plant (Figure 3c, Supplementary Figure 3a). This was contrary to the transcriptional responses in microbes, which were increased by the presence of APO (Figure 2a,b). To understand which biological processes (BP) were attenuated by APO in response to bacterial inoculums, we performed hierarchical clustering on the normalised z-scores, which revealed three distinct clusters (Supplementary Figure 3b). Subsequent gene ontology (GO) term enrichment analysis revealed that all three clusters were enriched for BP terms linked to biotic stress responses, i.e., “defence response to fungus” (Cluster I), “response to wounding” (Cluster I and II), or “defence response to bacterium” (Cluster III) (Supplementary Figure 3c).

### Bacteria trigger few long-term immune responses

Given that the presence of bacteria is sensed by the plant and elicits a response from the immune system, we asked how immunity genes responded to the bacterial inoculums in mock vs. APO treatments. As reference, we used a set of 64 PTI- and 238 ETI-associated genes known to be differentially expressed upon pathogen sensing (Bjornson et al., 2021; Ngou et al., 2021). As expected, plant growth-inhibiting LjRoot171 elicited the strongest immune responses, including the upregulation of nine LRR genes (Supplementary Figure 3d,e). Still, the immune response was weak for all inoculations, indicative of a return to steady-state transcription at the sampling point (3 weeks after inoculation) (Supplementary Figure 3d,e).

### JA-regulated chemical defence arsenal production depends on the chemical environment

Given the late sampling time point, we expected regulation of genes coordinating long term association of microbes with plants. Jasmonic acid (JA) is one of the main plant hormones involved in the regulation of plant defences against biotic stress, including the synthesis of defensive compounds (Pangesti et al., 2016; Zhang et al., 2021) and cross-talk with other hormones (Jang et al., 2020). To investigate the involvement of JA in the responses to bacterial inoculums, we interrogated the expression of genes involved in JA biosynthesis (Figure 3d) and downstream JA signalling (Figure 3e). In mock conditions, inoculation of bacteria led to a strong upregulation of *LOX2*, encoding for a lipoxygenase essential for JA biosynthesis (Mochizuki et al., 2016). In APO, the upregulation was around five-fold weaker (Figure 3d). The response of other JA biosynthesis genes (*AOS*, *AOC3* and *OPR3*) to single-strain inoculums were similarly dampened in APO compared to the mock environment (Figure 3d). Additionally, the JA-regulators *JAZ1, JAZ5* and *JAZ10* were upregulated upon inoculation in mock versus slightly downregulated in APO (Figure 3e). As a master regulator of plant defence responses, JA regulates the production of several plant specialised metabolites. We investigated the expression of genes from two major defence metabolite biosynthesis pathways downstream of JA, glucosinolates and flavonoids (Figure 3f-h, Supplementary Figure 4a,b). As expected, several of these genes were upregulated in bacteria-inoculated samples compared to axenic samples in mock conditions (Figure 3f-h; Supplementary Figure 4a,b). Among the glucosinolate biosynthesis genes, the effect was stronger for genes involved in the biosynthesis of aliphatic than for indolic glucosinolates (Figure 3h, Supplementary Figure 4a), suggesting a preferential production of aliphatic glucosinolates. By contrast, inoculated APO-treated samples showed a strongly reduced expression of genes involved in both glucosinolate and flavonoid synthesis, compared to the respective axenic control (Figure 3f-h, Supplementary Figure 4a,b). Taken together, these results indicate that the presence of APO leads to a dampening of the transcriptional activity of root chemical defence genes by interfering with JA synthesis and signalling.

### APO modulates sugar transport

Plants exert chemical control over their microbiomes through the production of defence compounds, but also by regulating nutrient availability, with sugars as major nutrients involved in plant-bacteria interactions (Chandran, 2015; Chen et al., 2010). We investigated the expression of two families of sugar transporters known to play a role in nutrient exchange during root colonisation (Chen et al., 2023), SWEETs and MSTs (monosaccharide transporters). Several *SWEETs* showed differential expression depending on the inoculum, with *SWEET3*, *SWEET11* and *SWEET12* down-regulated in presence of LjRoot171, and *SWEET13* and *SWEET14* up-regulated upon MF196 inoculation, independent of chemical treatment (Figure 3i). This indicated that APO did not influence the expression of these sugar transporters. MSTs behaved differently from SWEETs: *STP8*, *PLT3* and *ERD6L-14* were upregulated in mock but not in APO (Figure 3i), reminiscent of what we observed for genes coding for specialised metabolite biosynthesis. We thus concluded that the non-responsiveness of defence-related genes upon inoculation in the presence of APO is a broader phenomenon that targets both production of defence compounds as well as nutritional responses.

### Micro-communities elicit distinct response from their individual members

In natural conditions, plants are surrounded by several different bacteria attempting to colonise the root simultaneously. We looked at plant responses when inoculated with micro-communities, i.e., the pairs Root179xLjRoot171 and Root179xMF196. Interestingly, the micro-communities elicited much stronger responses than the single inoculums (Table 1, Figure 4a, Supplementary Figure 2a, Supplementary Figure 5a).

**Figure 4.**
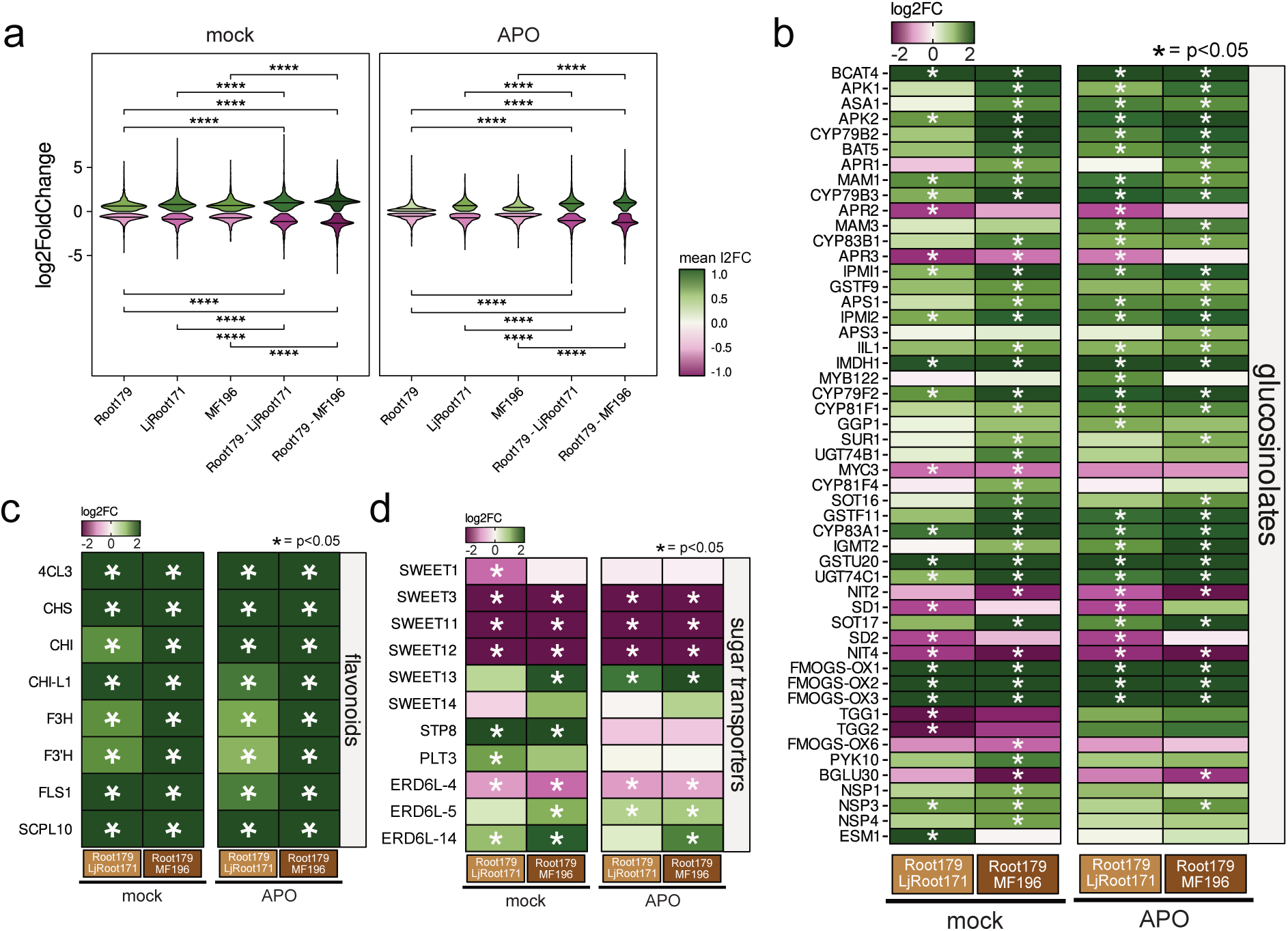
Micro-communities rescue APO-mediated dampening of defence responses. **a.** L2FC of all DEGs in samples inoculated with single isolates and micro-communities, treated with mock (left) or APO (right). ****: p-value < 0.0001 (two-tailed Student’s t-test). **b, c.** L2FC of genes involved in the biosynthesis of the defence compounds from the glucosinolate (b) and flavonoid (c) families showing differential expression in at least one condition. **d.** L2FC of genes encoding for sugar transporters that showed differential expression in at least one condition. Asterisks in b, c and d indicate significant differential expression (FDR < 0.05).

**Table 1.**
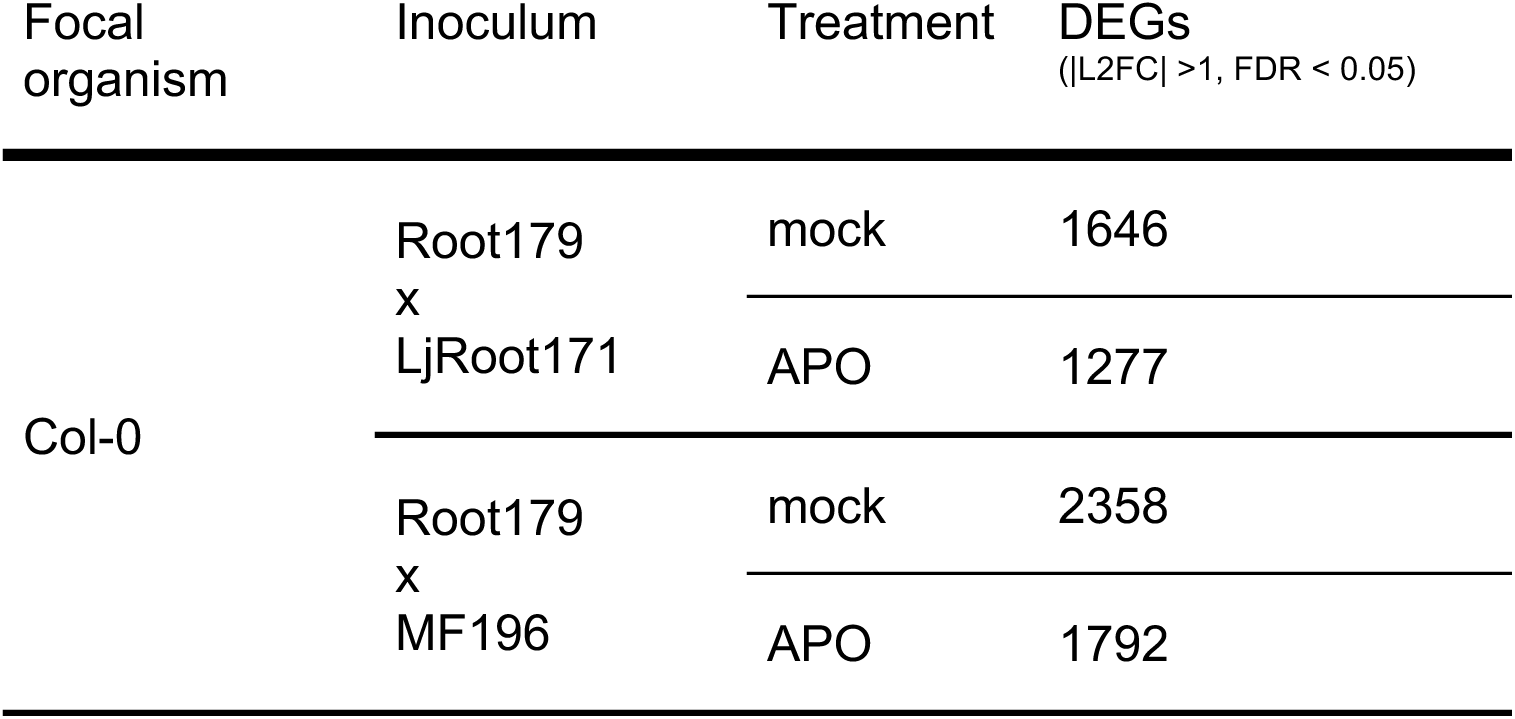
Effects of micro-community inoculums on the host plant. Col-0 differentially expressed genes (DEG, |L2FC| >1, FDR < 0.05) when inoculated with micro-communities, comparing the transcriptomics responses of axenic plants to their inoculated counterpart.

A PCA on the 1,000 Col-0 genes with the highest variance showed a separation of both micro-communities from their single-inoculum counterparts (Figure 3a). The samples from the Root179xLjRoot171 micro-community clustered closer to the LjRoot171 single-inoculum than to the Root179 single-inoculum samples, suggesting that LjRoot171 was the major driver of plant responses in this micro-community. The micro-community of Root179xMF196, showed a clearer separation from the axenic samples than the respective single inoculums. This grouping was also confirmed through hierarchical clustering based on normalised z-scores (Supplementary Figure 5b). Altogether, these results indicated that the responses elicited by micro-communities are not just the sum of the responses elicited by their respective single inoculums but involve more complex regulations of the members of these communities.

### Micro-communities rescue suppression of defence responses by APO

Focusing on genes involved in the biosynthesis of plant defence chemicals that were differentially regulated in response to APO, we checked their transcriptional response to both micro-communities. Surprisingly, the dampening of transcriptional response in APO compared to mock that we observed for single-strain inoculums (Figure 3f-h) was not detectable in response to micro-communities (Figure 4b,c, Supplementary Figure 6a,b). Instead, the flavonoid and glucosinolate pathways showed a strong and consistent upregulation independent of the chemical environment. Moreover, in Root179xMF196 micro-community inoculums, indolic glucosinolate biosynthetic genes, which had not responded to single isolates, were upregulated regardless of the chemical treatment.

To better understand what was driving these community-specific responses (Supplementary Figures 3a, 5a), we performed a PCA on the normalised counts of Col-0 genes when inoculated with micro-communities only (Supplementary Figure 7 a,b). Mock and APO treatments clearly separated, indicating an influence of the chemical environment on Col-0 genes in micro-communities. This was in contrast to the limited influence APO had on plant responses to single inoculums (Figure 3a). Interestingly, the single gene mostly driving the clustering by treatment in the Root179xLjRoot171-inoculated samples (Supplementary Figure 7a) encoded for an oxygenase responsible for the hydroxylation of JA to 12-OH-JA (*AT2G38240*; Caarls et al., 2017), important in the activation of the JA signalling pathway. This gene was strongly upregulated (l2FC = 4.8) in APO compared to mock in Root179xLjRoot171-inoculated roots. This suggested an increased defence response in APO in micro-communities compared to single inoculums, which might be linked to an activation of the JA signalling pathway via this oxygenase. Similarly, in the Root179xMF196 inoculated samples (Supplementary Figure 7b), the genes *BT1* and *BT2* (*BTB* [*broad-complex, tramtrack and bric-a-brac*]-*TAZ* [*transcriptional adapter zinc finger*]-*domain containing proteins 1 and 2*) showed strong contribution to the clustering by treatment. Genes from the BT family are also positively regulating the JA signalling pathway (Du et al., 2024; Zhou et al., 2022). Consistently, genes involved in JA biosynthesis and signalling pathways were upregulated in response to micro-communities (Supplementary Figure 7c,d). In single-strain inoculums, APO led to the dampening of the upregulation of these pathways (Figure 3 d,e). In micro-community inoculums, we observed the opposite trend (albeit not statistically significant) (Supplementary Figure 7c,d). We concluded that bacterial communities can counteract the chemical interference on the JA pathway, leading to increased plant defence responses in presence of bacterial communities. Surprisingly, *SD1* (*Sulfur deficiency-induced*, *AT5G48850*) also showed strong contribution to the separation by treatment (Supplementary Figure 7b) and was upregulated in APO in Root179xMF196-inoculated plants (Figure 4b). This gene has been shown to downregulate aliphatic glucosinolate biosynthesis (Aarabi et al., 2016), a response that we only observed in APO when plants were inoculated with single isolates, suggesting that other mechanisms might be regulating glucosinolate biosynthesis in micro-community-inoculated plants.

## DISCUSSION

### Bottom-up and top-down approaches converge to unravel chemical influences on bacterial interactions

The establishment of a specific microbiome on the host roots is a tightly regulated process that can be influenced by external factors (Trivedi et al., 2020). Specialized metabolites are secreted by plants into the soil, impacting the microbiome establishment of current and future hosts (Gfeller et al., 2023; Steinauer et al., 2023; Stengele et al., 2024). In this study, we used a bottom-up approach to initially study the impact of the chemical environment on bacteria-bacteria interactions. In our large-scale binary interaction assay, we found that the benzoxazinoid-derived compound APO and its precursor BOA can both influence bacteria-bacteria interactions via processes that are independent from the direct effects of the chemicals on bacterial growth.

The subsequent top-down approach to decipher the mechanistic impact of APO on plant-microbe interactions indicated that such altered interactions can, in at least one case, be traced back to resource divestments towards detoxification, leading to decreased competitiveness. The decreased competitive ability of Root179 towards LjRoot171 in the presence of APO was linked to an upregulation of pathways associated with detoxification, and a cognate downregulation of the offensive arsenal, such as the type-II secretion system (TIISS) in Root179. We propose that this decreased competitive ability resulted in an APO-specific inhibition of LjRoot171 towards Root179. Overall, our findings highlight the relevance of the chemical dimension in the bottom-up design of communities. Accounting for the chemical environment and potential fluxes is an important design element to ensure that observed interactions that contribute to the ecosystem, for example between keystone and other taxa, are robust to chemical perturbations.

### The host’s ability to sense microbes is modulated by the chemical environment

Recently, Ordon et al. (2025) highlighted the importance of the Root179 TIISS in the context of host colonisation, with two type-II secreted enzymes suppressing the sensing of Root179 by Col-0. In line with these results, we found that different isolates lead to different transcriptomic responses in Arabidopsis. While the plant mounted a strong immune response to the pathogenic isolate LjRoot171, only a diminished response was observed to the commensal bacteria Root179 and MF196. Notably, the presence of APO resulted in a weaker response to LjRoot171. Taken together, these results indicate a complex signal integration, where the response to a micro-community of bacteria can be balanced by the chemical environment.

### Chemical defences are dampened by APO and amplified by micro-communities

Micro-communities elicited stronger responses at the level of secondary metabolism than single inoculums, in particular in the glucosinolate and flavonoid biosynthesis pathways. The observation that the biosynthesis pathways of these two main defence compounds were impaired further pointed towards APO inhibiting the ability of the plant to respond to colonizing bacteria. We also observed a more subtle differentiation in the response of the glucosinolate pathway. Single-strain inoculums upregulated mostly genes involved in the synthesis of aliphatic and lesser in indolic glucosinolates in mock-treated plants, suggesting a targeted plant response with preferential production of one type of glucosinolates. In contrast, both pathways were upregulated upon colonisation by micro-communities. Genes belonging to the flavonoid biosynthesis pathway were differentially expressed between the two chemical environments in single inoculums. Notably, chalcone synthase (CHS) and chalcone isomerase (CHI), catalysing the synthesis of naringenin and thus the first dedicated steps in the flavonoid/anthocyanin pathway (W. Liu et al., 2021) were consistently upregulated in single inoculums in mock conditions. Conversely, micro-communities strongly induced the expression of the flavonoid but not the anthocyanin pathway, indicating that the plant mounts a stronger and more specialised metabolic response when facing multiple bacteria, independent of the chemical environment (Figures 3,4). Overall, we conclude that with increasing complexity of the biotic environment, the influence of the chemical environment weakens; signal integration by the host that underlies the output response hence appears to become less sensitive to a single chemical compound if the biotic environment increases in complexity.

### Struggle for sugar

Carbon provision by the host towards colonizing microbes is one of the major drivers of host colonisation (Estabrook and Yoder, 1998). In both single inoculums and micro-communities, we observed a complex regulatory landscape of sugar exporters, modulated by the chemical environment. The host plant responded with an upregulation of *SWEET13*, and to a lesser extent *SWEET14*, in response to *Bacillus* MF196. By contrast, inoculums with the growth-inhibiting strain LjRoot171 resulted in a downregulation of *SWEETs 11*, *12* and *3*. The response to Root179, previously shown to be able to camouflage itself from host recognition (Ordon et al., 2025), did not lead to a consistent differential expression of *SWEETs;* the downregulation of *SWEET3* in mock conditions did not occur in presence of APO. As SWEETs are canonically mediating the efflux of sugars into the apoplastic space (Chen et al., 2010), we conclude that the host attempts to restrict pathogens by decreasing sugar availability.

Notably, when comparing expression of SWEETs between single inoculums and micro-communities, both micro-communities elicited a SWEET response similar to that of LjRoot171, with a downregulation of *SWEETs 11*, *12* and *3*; even though the ability of MF196 ability to increase expression of *SWEET14* remained, even when embedded in a micro-community. SWEETs, and particularly SWEET-11 and -12, were previously shown to play a role in the longitudinal organisation of the microbiome along the root axis (Loo et al., 2024). The downregulation of the expression of both of those transporters in response to bacteria inoculums further points towards an active modulation of root colonisation by the host.

Interestingly, expression of *STP8*, a member of the major facilitator superfamily of transporters and a transporter of hexoses (J. Liu et al., 2021), was impacted by APO in the presence of bacteria. *STP8* was upregulated in response to bacterial colonisation only in mock conditions, but not in the presence of APO. STPs are generally assumed to act antagonistically to SWEETs, transporting sugar from the apoplast into the intracellular space (Chen et al., 2024; Rottmann et al., 2018) and STP13 has previously been reported as responsive to flg22 and contributing to control of bacterial infection (Yamada et al., 2016). As unspecific glycosylation of APO has previously been suggested to be a route towards detoxification (Schulz et al., 2016), draining sugars from the apoplastic space might be unfavourable in the presence of APO, as the host tries to balance chemical stress and colonisation by microbes.

## CONCLUSION

Taken together, our study highlights the role of legacy metabolites on root colonizing communities. We showed that dissecting higher-order interactions to the lowest levels of complexity before reconstructing more complex communities is a key consideration when setting out to understand the fine-tuning effect of soil legacy on plant-microbe interactions, emphasising the importance of decrypting lower-level interactions to understand the bigger picture. In particular, the dampening of plant defence responses by long-lasting soil chemicals might be relevant in an agricultural context. Future research should take into account residual metabolites in soil in addition to existing soil communities, to adequately reflect the environment of plants, and specifically agricultural crops.

## MATERIAL AND METHODS

### Bacterial isolates sequencing and cultivation

Bacterial collections were obtained from Paul Schulze-Lefert (Max Plant Institute for Plant breeding Research, Cologne, Germany) and Jeffrey Dangl (University of North Carolina at Chapel Hill, Michigan, USA). The identity of all bacterial isolates used in this study was confirmed via Illumina whole-genome sequencing. Genomic DNA was extracted using the Monarch Genomic DNA purification kit (New England Biolabs); indexed libraries were prepared according to the protocol of Jones et al. (2023). Briefly, the DNA was fragmented (3.6 ng DNA, 0.45 µl TDE1 enzyme (Illumina #20034197), 5 µl 2x tagmentation buffer, water to 10 µl) at 55 °C for 10 min. The barcoding PCR was performed with indexed primers (Jones et al., 2023: Supplementary material S2) fitting the TDE1 adaptors and Q5 high-fidelity DNA polymerase (New England Biolabs, #M091S), with the corresponding buffer and high-GC enhancer, in a total reaction volume of 17 µl. The PCR reaction was as follows: 72 °C - 3 min; 98 °C - 30sec; [98 °C - 10 sec; 63 °C - 30 sec; 72 °C - 1 min] x 10; 10 °C - hold. 6 µl of each library were pooled together, and were cleaned and size selected for fragments of 400bp on average using MagSi-NGS PrepPlus magnetic beads (Magtivio, ratio 0.8). Paired-end Illumina sequencing was carried out by Novogene UK (Cambridge, United Kingdom) with 500 Mb per sample (2×150 bp paired-end on an Illumina NovaSeq X Plus instrument). Reads were mapped to the reference genomes (http://www.at-sphere.com/, http://labs.bio.unc.edu/dangl/) in a custom Nextflow pipeline and further analysed using custom R scripts. In brief, reads for each isolate were mapped against the reference genome using minimap2 (Li, 2018). Subsequently, unmapped reads were mapped against all other genomes of strains in the collection, to identify cross-contamination events. Isolates with more than 80% of reads mapping to the expected genome were classified as confirmed. Out of 237 obtained isolates, 73 could be confirmed by whole-genome sequencing. Isolates where the identity could not be confirmed were excluded from the study.

The focus isolates from this study were re-sequenced to obtain high-quality genomes. The genome of Root179 was sequenced using Oxford Nanopore Technology (PromethION 24), genomes of LjRoot171 and MF196 were assembled from PacBio CLR reads (R. Garrido-Oter, unpublished) and polished using paired-end Ilumina sequencing reads, carried out by Novogene GmBH (Martinsried, Germany), (2×150 bp paired-end) on an Illumina NovaSeq X Plus instrument.

Assembly and annotation were performed using a custom Nextflow pipeline (https://gitlab.lrz.de/beckerlab/nf-bacassemblotate), using Flye (v2.9.2; Kolmogorov et al., 2019) for assembly, Pilon (v1.24; Walker et al., 2014) for polishing and BAKTA (v1.9.3; Schwengers et al., 2021) for annotation.

Unless stated otherwise, the isolates were inoculated from the glycerol stock on ½ TSB agar media (8.5 g casein hydrolysate, 1.25 g D-glucose, 1.25 g di-potassium hydrogen phosphate, 1.5 g soy peptone, 2.5 g sodium chloride, 15 g bacto agar, dissolved in 1 L water and pH adjusted to 6.8) and grown at 28°C for 1-3 d.

### Bacterial collection screening

To assess APO production, one loop of culture was inoculated onto a fresh plate of ½ TSB agar supplemented with 500 µM BOA or the same volume of the solvent DMSO as a control, and incubated at 28°C for 10 d. After 10 d, APO production was assayed by visually detecting red colour change on the BOA plates. APO sensitivity was assessed by growing the bacterial isolates in media supplemented with APO to a final concentration of 50 µM, in a 96-well plate assay described previously (Schandry et al., 2021; Thoenen et al., 2023). Briefly, each isolate was inoculated from a pre-culture into a 96-well plate, with 4 replicates per isolate and per condition. The plates were incubated in an automated plate reader (Tecan Spark, TECAN) and the absorbance at 600 nm was read every 20 min for 3 d. The area under the curve (AUC) was calculated from the growth curves, and treatment AUC was compared to the AUC of the control condition. Isolates with a relative AUC lower than 50% of the control were marked as sensitive.

### Pairwise interaction assay

All bacterial isolates were tested for their inhibition of other isolates, using a modified protocol from Getzke et al. (2023). Briefly, isolates were grown in 1 ml ½ TSB overnight, and 100 µl of culture was mixed with 60 ml molten ½ R2A agar (Merck) at 45°C, then poured into a square plate. 10 µl of overnight cultures of the isolates to be tested for inhibition was inoculated in 200 µl ½ TSB media 96-well plates and shaken for 1 h at 28°C. A drop (around 5 µl) of the dilution was plated on the solidified bacteria-containing agar using a replicator and the plates were incubated at 22°C. Presence of inhibition halos were marked after 5 d of incubation. The assay was repeated once with independent biological replicates.

To test the effects of benzoxazinoids, BOA, APO or DMSO (solvent-control, ‘mock’) were added to the molten agar prior to pouring the plates, to a final concentration of 500 µM (BOA), 50 µM (APO) or the same volume of DMSO.

### Plant growth conditions in calcined clay

Plants were cultivated as described in Schäfer et al. (2022), with some modifications. *Arabidopsis thaliana* Columbia-0 (Col-0) seeds were surface-sterilised as above. The wells of 6-well plates were filled with 5 ml of calcined clay (drying agent, DiamondPro), previously washed and twice-autoclaved. 2 ml of 1/2MS + vitamins (glycine 0.002 g/L; myoinositol 0.1 g/L; nicotinic acid 0.0005 g/L; pyroxidine-HCl 0.0005 g/L; thiamine-HCl 0.0001 g/L) was added to each well. Individual seeds were placed in each well and the plates were sealed with micropore tape. The same number of seeds was plated on 1/2MS agar plates. All plates were placed in the dark at 4°C. After 4 d, the plates were transferred to the growth cabinet (same conditions as above). After 6 d, each well was watered with 200 µl of 1/2MS + vitamins and seedlings from the agar plates were transferred in the wells that showed no visible seedling growth. After 8 d, the seedlings were inoculated with the corresponding bacteria and chemical treatment (see below).

### Inoculation with bacteria and chemical treatment

Bacterial isolates were cultivated, washed and resuspended in MgCl_2_ as described above. The washed cultures were diluted to an optical density at 600 nm (OD_600nm_) of

0.02 and 200 µl (for single isolate inoculation) or a mix of 100 µl of each isolate (for dual inoculation) was inoculated at the base of the seedling. For benzoxazinoid inoculation, 200 µl of a 500 µM (BOA), 50 µM (APO) or DMSO (mock) solution was added at the base of the seedling. The lids were sealed to the plates using micropore tape and incubated for 20 d in a growth cabinet using the same conditions as above. Each well was watered twice a week with 200 µl 1/2MS + vitamins.

### Plant harvesting and CFU count

After 28 d, pictures were taken from each plate with a camera at a fixed distance from the top of the plate. Using sterile tweezers and razor blade, the roots were detached from as much clay as possible, rinsed in 1ml sterile water, separated from the leaves, placed in an Eppendorf tube, flash-frozen in liquid nitrogen and stored at −70°C. For CFU count, 100 µl of the water used for rinsing was transferred to a 96-well plate, and an 8-step 10-fold dilution series was performed. 10 µl from each dilution step was plated on TSB agar, the plates incubated at 28°C, and the colonies were counted after 1-2 d of incubation.

### Total RNA extraction and sequencing

Total RNA was extracted using a CTAB method (Bemm et al., 2016). Briefly, the roots of 3 biological replicates were pooled and manually ground in liquid nitrogen. 700 µl of hot (65 °C) RNA extraction buffer (2 g CTAB; 2 g polyvinylpyrrolidone 40 g/mol (PVP40); 11.7 g NaCl; 10 ml TrisHCl 1M; 5 ml Na-EDTA 0.5M; 100 ml DEPC water), 7 µl of Tris-Carboxyethylphosphine (TCEP) and one small spatula of polyvinylpolypyrrolidone was added to each tube and mixed thoroughly. The samples were incubated at 65°C, 600 rpm for 10 min. 700 µl of chloroform:isoamylalcohol (24:1) was added to each tube and mixed by inversion. The tubes were centrifuged for 8 min at 10.000 rcf, the upper phase was transferred to a new tube, 175 µl lithium chloride (8 M LiCl) was added to each tube and the RNA was precipitated overnight at 4°C. The next day, the precipitated RNA was pelleted in a 4°C pre-cooled centrifuge at 20.000 rcf for 20 min and resuspended in 100 µl DEPC water. 10 µl sodium acetate (3 M NaOAC) and 250 µl 96% ethanol was added to each tube, and frozen at −20°C for 1 h. The precipitated RNA was pelleted in a pre-cooled centrifuge as above, washed with 500 µl 80% ethanol, centrifuged at 4°C, 20.000 rcf for 20 min. The pellet was briefly dried at 37°C and resuspended in 20 µl DEPC water. RNA concentration was measured on a NanoDrop (ThermoScientific) and the quality was assessed on a BioAnalyzer (Agilent Technologies). Total RNA was treated with DNase (New England Biolabs) according to the manufacturer’s instructions. Dual rRNA depletion and paired-end sequencing was carried out by Novogene GmBH (Martinsried, Germany) with >100 million reads per sample (2×150 bp paired-end on an Illumina NovaSeq X Plus instrument).

### Data analysis

The raw data was processed using the nf-core/rnaseq (v3.14.0; Patel et al., 2024). The reads were mapped to the Col-0-CEN reference genome (Naish et al., 2021) and to the *de-novo* assembled bacterial genomes. After filtering low quality reads, we could unambiguously map 33-83 million reads per sample (Supplementary Figure 1d). A negligible part (<1%) of all filtered reads could not be mapped to the provided references, validating that the samples were free of contamination. We removed transcripts with a read count <10 and performed comparative analysis per organism using DESeq2 (v3.20, Love et al., 2014).

All scripts and data used in this study can be found on the GitHub repository https://github.com/lrouyer/Rouyer_et_al_interactions.git.

## DATA AVAILABILITY

The files and code used for the analysis will be made available on https://github.com/lrouyer/Rouyer_et_al_interactions.git upon final publication of the manuscript. Sequencing data will be deposited in the European Nucleotide Archive (ENA; ebi.ac.uk/ena) under the project number PRJEB90180 upon final publication of the manuscript.

## ACKNOWLEDGEMENTS

We would like to thank Silke Robatzek, Duncan Crosbie and Alessa Ruf for their critical reading of the manuscript and constructive remarks. Thank you to Paul Schulze-Lefert (MPI-PZ Cologne) and Jeffery Dangl (UNC Chapel Hill, NC) for sharing their bacterial collections. We would like to also thank Ruben Garrido-Oter (MPI-PZ Cologne) for giving us access to long-read data from the AtSphere and LjSphere bacterial collection, and Stefan Krebs (Gene Center, LMU Munich) for the support in bacteria whole genome sequencing. We are grateful to the Life Science Munich Graduate program for their support of L.R., and to Philipp Chapman, Pubali Paul and Sylvia Devkota for their valuable technical assistance.

## FUNDING

This work was supported by the European Union’s Horizon 2020 research and innovation programme by the European Research Council (ERC) (Grant Agreement No. 716823 ‘FEAR-SAP’ to C.B.); and by the Deutsche Forschungsgemeinschaft (DFG) in the frame of TRR356 project A05 (project no. 491090170 to N.S.) and in the frame of the SPP2125 ‘DECRyPT’ (project no. 466385132 to N.S. and C.B.). For the computational analyses, we used the BioHPC-Genomics compute cluster at the Leibniz Rechenzentrum Munich (DFG; grant no. 416 450674345) and used infrastructure provided by TRR356 project I01.

## AUTHOR CONTRIBUTIONS

L.R., N.S. and C.B. designed the research; L.R. performed the experiments and analysed the data; and L.R., N.S. and C.B wrote the paper.

## CONFLICT OF INTEREST

The authors state that they do not have a conflict of interest.

## SUPPLEMENTARY MATERIAL

Supplementary Figure 1. Experimental design and validation of meta-transcriptomics dataset.

Supplementary figure 2. Transcriptomic responses of bacteria to co-inoculation in micro-communities.

Supplementary figure 3. Transcriptional response of Arabidopsis to single bacteria inoculation and immunity genes expression.

Supplementary figure 4. Transcriptional response of glucosinolate and flavonoid associated genes to single bacteria inoculation.

Supplementary figure 5. Transcriptional response of Arabidopsis to micro-community inoculation and hierarchical clustering of samples.

Supplementary figure 6. Transcriptional response of glucosinolate and flavonoid associated genes to micro-community inoculation.

Supplementary figure 7. Responses of defence and JA-associated Arabidopsis genes to micro-community inoculation.

## Notes

### Competing Interest Statement

The authors have declared no competing interest.

