## Supplementary figures for "Plant specialised metabolites modulate the molecular signatures of host-bacteria and bacteria-bacteria interactions"

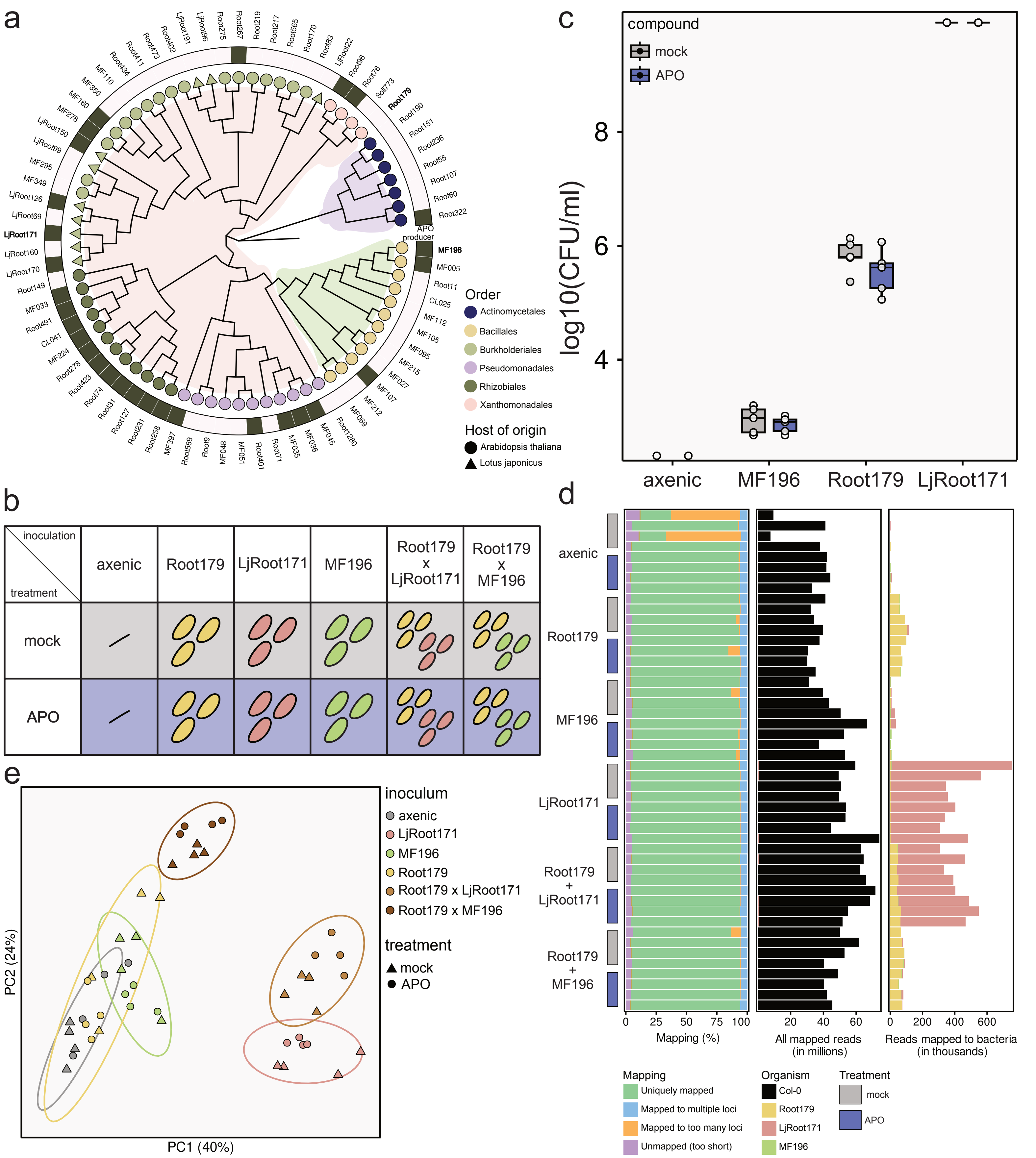

**Supplementary figure 1. Experimental design and validation of meta-transcriptomics dataset. a.** Phylogenetic tree of bacteria used in this study. Outer ring indicate bacteria that are able to convert BOA to APO. **b.** Experimental design of the meta-transcriptomics samples. **c.** Log10 transformation of colony forming units (CFU) counts per ml of roots inoculated with different bacteria and treated with DMSO (mock) or APO. **d.** Mapping rates of all reads (left), Col-0 reads (middle) and bacteria reads (right) of total RNA extracted from 4-week old Arabidopsis thaliana inoculated with different bacteria and treated with control or APO. **e.** Principal component analysis (PCA) of all reads from the meta-transcriptomics analysis.

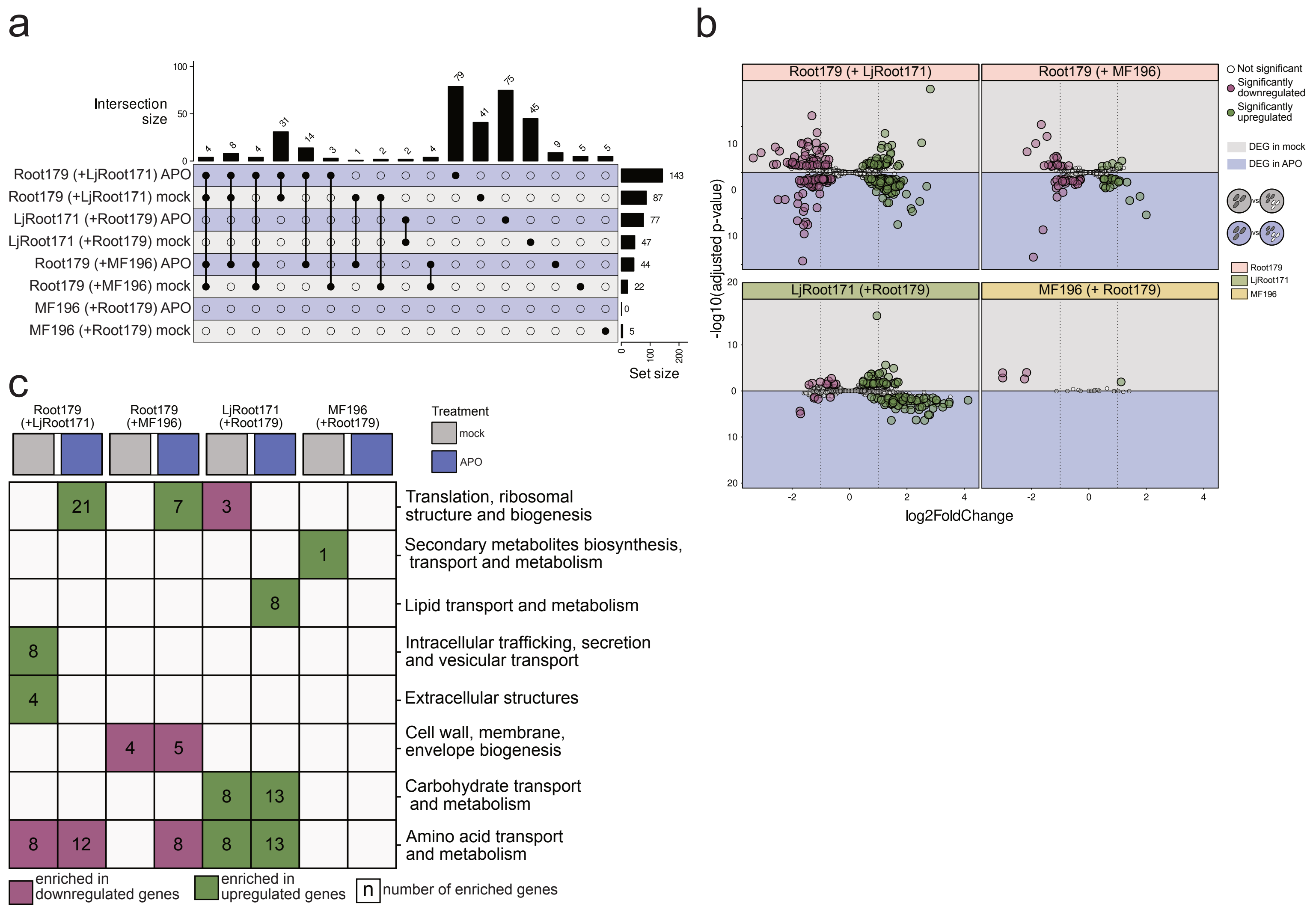


**Supplementary figure 2. Transcriptomic responses of bacteria to co-inoculation in micro-communities. a.** Upset plot of bacteria DEG with |log2FC| > 1 per inoculum and overlapping DEGs when comparing bacteria alone and in micro-communities. **b.** Volcano plot of differentially expressed genes in bacteria co-inoculated with other bacteria and treated with mock (grey background) or APO (purple background). **c.** Cluster of orthologuous genes (COG) term analysis of all differentially expresed genses in bacteria.

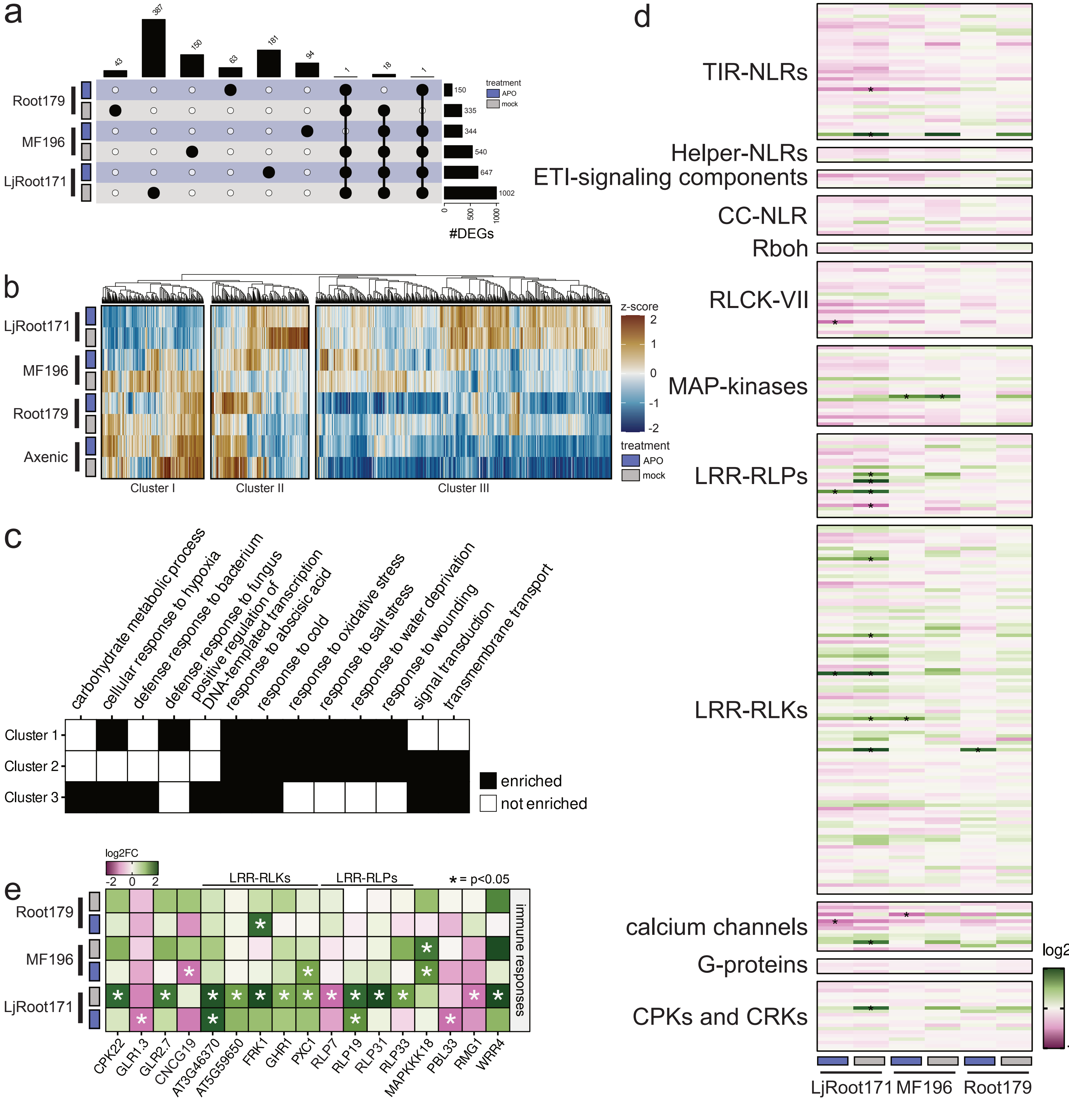


**Supplementary figure 3. Transcriptional response of Arabidopsis to single bacteria inoculation and immunity gene expression. a.** Upset plot of Arabidopsis DEG with |log2FC| > 1 per inoculum and overlapping DEGs when comparing axenic and bacteria-inoculated roots. **b.** Z-score hierarchical clustering of normalized counts of genes showing differential expression in bacteria-inoculated Col-0. **c.** Gene ontology (GO) term enrichment of genes present in Clusters 1, 2 and 3 of the z-score heatmap in panel a. **d.** Log2FC of immunity-related genes that showed differential expression in previous studies (Bjornson et al. 2021, Ngou et al. 2021). **e.** Log2FC of immunity-related genes that showed differential expression in our study in at least one condition. Stars indicate a FDR < 0.05.

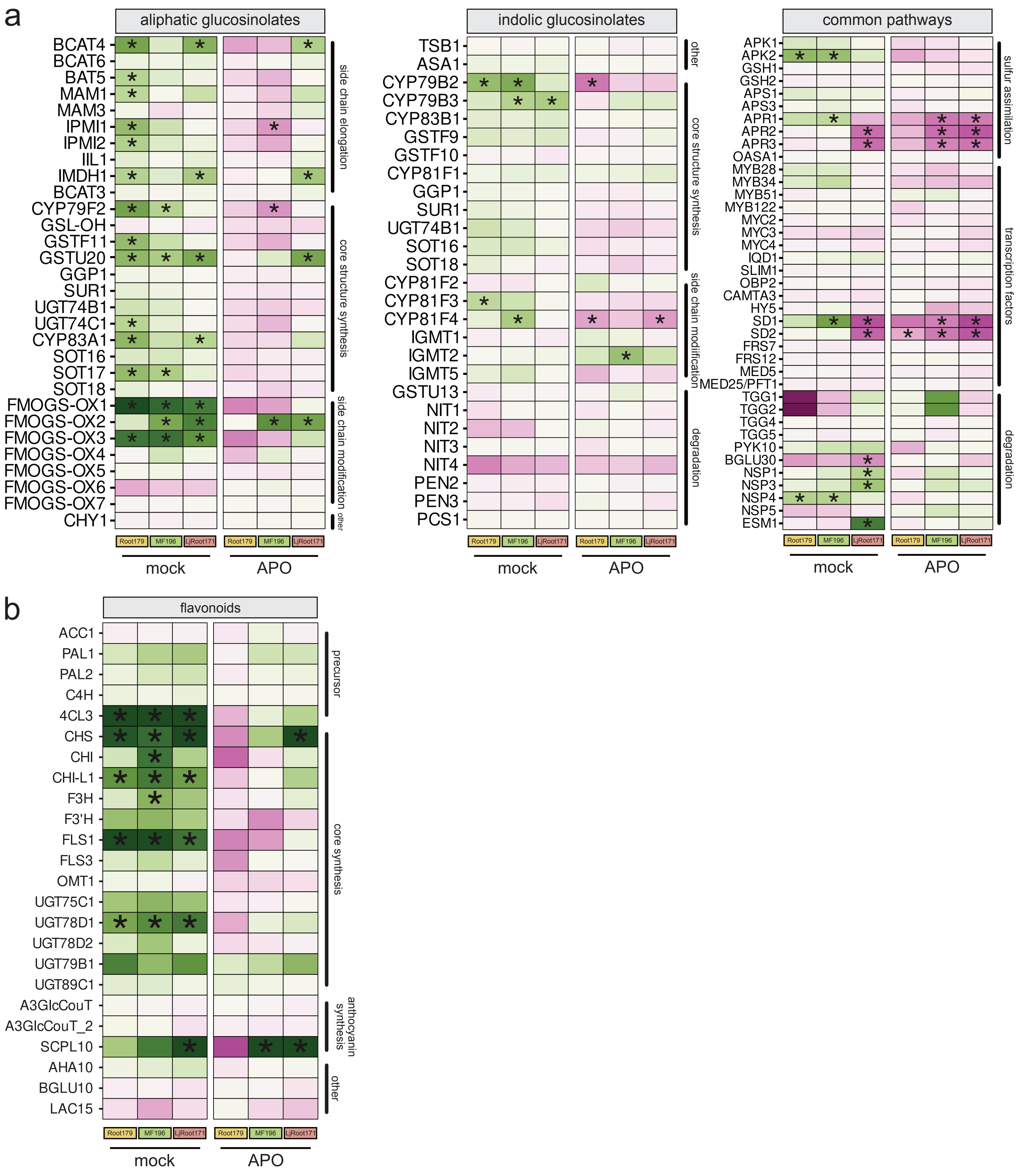


**Supplementary figure 4. Transcriptional response of glucosinolate and flavonoid associated genes to single bacteria inoculation. a.-b.** Log2FC of all genes involved in the synthesis of the defense compounds from the glucosinolate (a) and flavonoid (b) families detected in our dataset. Stars indicate a FDR < 0.05.

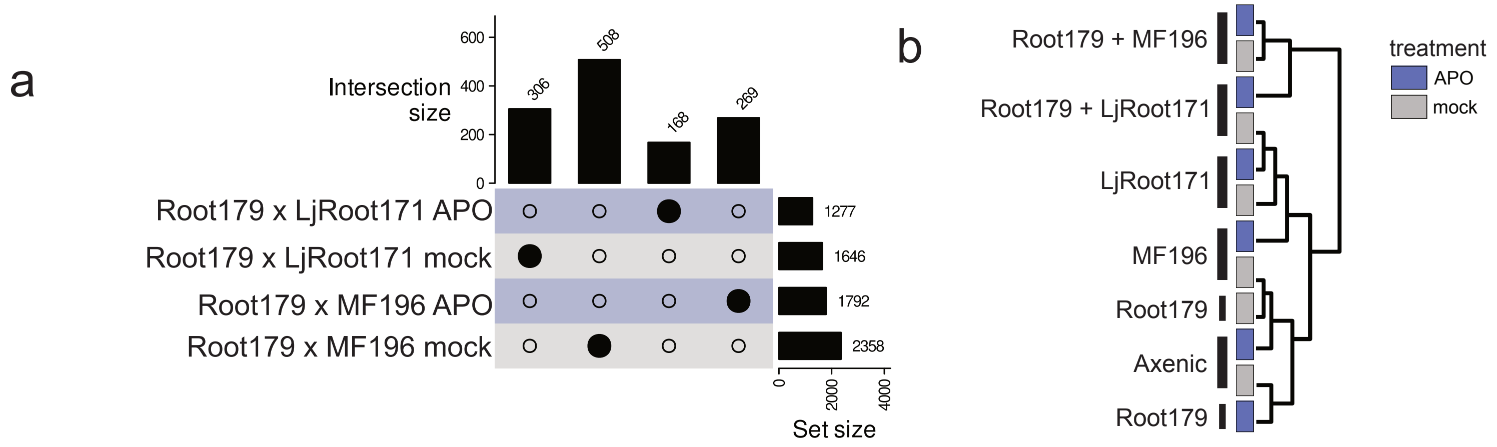


**Supplementary figure 5. Transcriptional response of Arabidopsis to micro-community inoculation and hierarchical clustering of samples. a.** Upset plot of Arabidopsis DEG with |log2FC| > 1 per inoculum and overlapping DEGs when comparing axenic and roots inoculated with micro-communities. **b.** Hierarchical clustering of samples based on the z-scores of normalized counts of Col-0 genes showing differential expression.

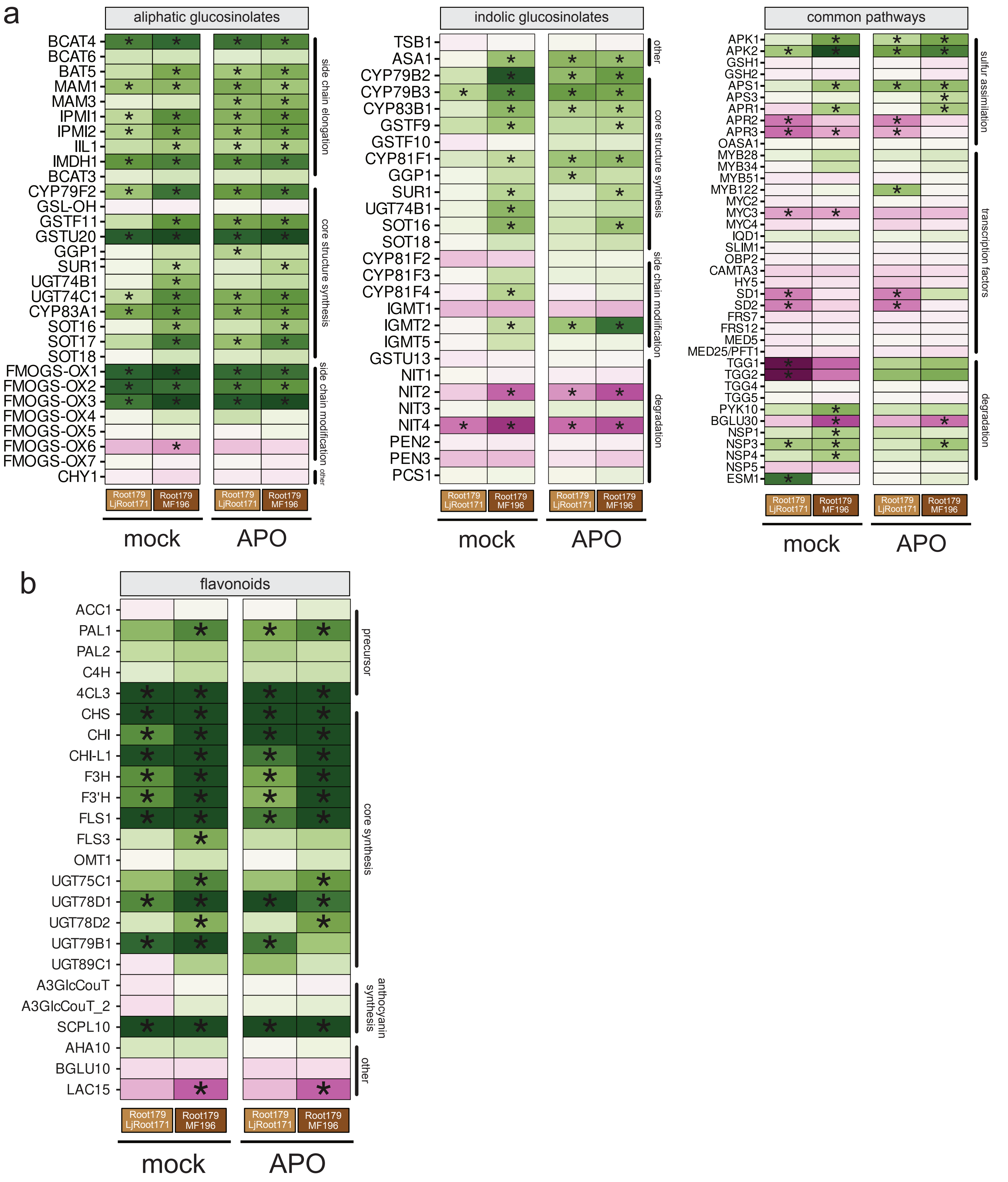


**Supplementary figure 6. Transcriptional response of glucosinolate and flavonoid associated genes to micro-community inoculation. a.-b.** Log2FC of all genes involved in the synthesis of the defense compounds from the glucosinolate (a) and flavonoid (b) families detected in our dataset.

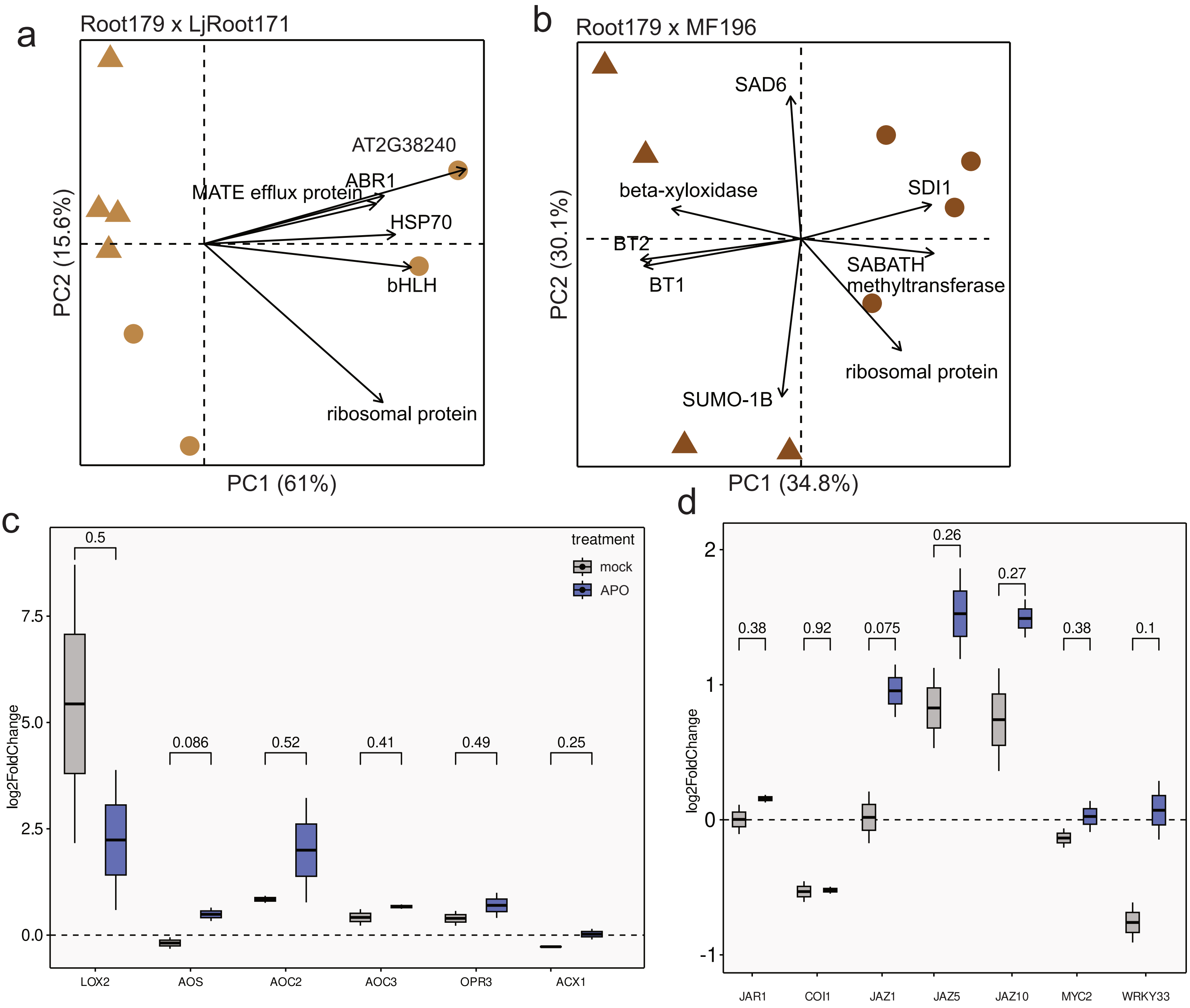


**Supplementary figure 7. Responses of defense and JA-associated Arabidopsis genes to micro-community inoculation. a,b.** Biplot of Col-0 genes showing the Root179xLjRoot171-inoculated samples (a) and the Root179xMF196-inoculated samples (b). Arrows represent the genes with the most important contribution to the sample separation, arrow length is proportional to the gene contribution. **c,d** Mean log2FC of genes involved in JA synthesis (c) and signaling (d).
